# Spindle-dependent memory consolidation in healthy adults: A meta-analysis

**DOI:** 10.1101/2022.07.18.500433

**Authors:** Deniz Kumral, Alina Matzerath, Rainer Leonhart, Monika Schönauer

## Abstract

Accumulating evidence suggests a central role for sleep spindles in the consolidation of new memories. However, no metaanalysis of the association between sleep spindles and memory performance has been conducted so far. Here, we report meta-analytical evidence for spindle-memory associations and investigate how multiple factors, including memory type, spindle type, spindle characteristics, and EEG topography affect this relationship. The literature search yielded 53 studies reporting 1427 effect sizes, resulting in a small to moderate effect for the average association. We further found that spindle-memory associations were significantly stronger for procedural memory than for declarative memory. Neither spindle types nor EEG scalp topography had an impact on the strength of the spindle-memory relation, but we observed a distinct functional role of global and fast sleep spindles, especially for procedural memory. We also found a moderation effect of spindle characteristics, with power showing the largest effect sizes. Collectively, our findings suggest that sleep spindles are involved in learning, thereby representing a general physiological mechanism for memory consolidation.

**Highlights:**

- Spindle measures showed a small to medium-sized association with memory performance.
- This relationship was stronger for procedural memory than declarative memory.
- No moderation effects of spindle type and EEG scalp topography have been observed.
- Spindle power emerged as the strongest predictors.
- Naps showed similar spindle-related consolidation mechanisms to whole-night sleep.

## 1. Introduction

A wealth of research, much of it conducted over the past 30 years, has shown that we retain memories better over a period of sleep relative to wakefulness. With the positive effect of sleep on memory being well-established, current research efforts have been directed at uncovering the mechanisms that mediate this relationship. Accumulating evidence suggests that sleep does not merely passively protect memories from interference and decay, but actively fosters their consolidation. Thus, longer sleep times lead to greater memory benefits (Diekelmann et al., 2012; Schönauer et al., 2014), the occurrence of sleep-specific brain oscillatory activity has been linked with memory consolidation (Fogel and Smith, 2006; Gais et al., 2002; Girardeau et al., 2009; Marshall et al., 2006; Perrault et al., 2019; Schmidt et al., 2006), and it has been discovered that learning-related brain activity is reactivated during non-rapid eye movement (NREM) sleep (Peigneux et al., 2003; Schönauer et al., 2017; Schreiner et al., 2021; Wilson and McNaughton, 1994). Sleep-dependent memory reactivation, in particular, is a promising candidate mechanism to underlie behavioral benefits of sleep on memory performance (Antony et al., 2012; Rasch et al., 2007; Rudoy et al., 2009). Memory reactivation has been linked to oscillations characteristic of NREM sleep, such as slow oscillations (Cousins et al., 2014; Ngo and Staresina, 2022; Schönauer et al., 2017; Schreiner et al., 2021), sleep spindles (Antony et al., 2018; Bergmann et al., 2012; Cairney et al., 2018; Fogel et al., 2017; Jegou et al., 2019; Laventure et al., 2018; Schönauer et al., 2017; Wang et al., 2019), and sharp-wave ripples (SWR) (Fernández-Ruiz et al., 2019; Rothschild et al., 2017; Zhang et al., 2018). These oscillations are coordinated within a network of brain regions including the thalamus, neocortex, and hippocampus. Research in both animals and humans has shown that the precise orchestration of these brain rhythms contributes to memory consolidation during sleep (Maingret et al., 2016; Mölle et al., 2009; Staresina et al., 2015). Sleep spindles, in particular, have emerged as a promising candidate marker for memory reactivation in the human brain and have been linked to behavioral benefits of sleep for memory. Spindle activity during sleep increases after an intense learning experience and is related to how well the memory is retained (Gais et al., 2002; Morin et al., 2008; Schabus et al., 2008), and artificially boosting sleep spindle occurrence by pharmacological interventions benefits memory consolidation (Mednick et al., 2013). Moreover, the presentation of learning-related sounds during sleep elicits stronger activity in the spindle frequency band than the presentation of control sounds (Cairney et al., 2018), and brain activity during sleep spindles is informative about the kind of learning material studied before sleep (Cairney et al., 2018; Schönauer et al., 2017). Because recent findings have shown that effect sizes for associations between sleep and memory performance have been overestimated (Ackermann et al., 2015; Berres and Erdfelder, 2021; Cordi and Rasch, 2021; Pöhlchen et al., 2021; Reverberi et al., 2020), we should quantitively evaluate the influence of specific sleep parameters on memory in a similar way. In this meta-analysis, we therefore investigated whether spindle activity shows a consistent link with memory performance, and overnight memory consolidation in particular.

### 1.1. The role of sleep spindles in memory consolidation and reprocessing

Sleep spindles are a physiological hallmark of NREM sleep, where they appear as brief oscillatory events with a burst-like sequence of 10–16 Hz sinusoidal cycles, lasting for at least 0.5 sec, but no longer than 2-3 sec (Fernandez and Lüthi, 2020; Lüthi, 2014). They are generated through interactions between inhibitory cells in the thalamic reticular nucleus and bursting thalamocortical relay neurons (Contreras and Steriade, 1996; Steriade, 2006). Cortical inputs to the thalamus may regulate spindle initiation and termination (Bonjean et al., 2011).Importantly, spindles are thought to play a major role in mediating neural plasticity (Peyrache and Seibt, 2020; Steriade and Timofeev, 2003) and memory replay (Siapas and Wilson, 1998).

A direct link between spindle activity and plasticity has first been proposed by Timofeev and colleagues (Timofeev et al., 2002), who showed that spindle activity is associated with long-term changes in responsiveness of cortical neurons. Similarly, input firing patterns during spindles have been found to induce long-term potentiation (LTP) effectively in neocortical pyramidal cells (Rosanova and Ulrich, 2005), probably by increasing intracellular calcium concentrations. Recent calcium imaging studies have also provided evidence that spindles may trigger cell- and layer-specific activation (Niethard et al., 2017), which could support plasticity underlying memory consolidation. It has been shown that the calcium activity (Ca2+) in neocortical dendrites is increased and synchronized during oscillations in the spindle range (Seibt et al., 2017), which may be enhanced through coupling with slow oscillation (SO) up-states (Niethard et al., 2018), thereby strengthening memory consolidation.

Sleep spindles are further thought to play a pivotal role in memory reactivation. Memory reactivation has mainly been studied in the rodent hippocampus, where it predominantly occurs during SWRs, which are transient high-frequency oscillations (140–200 Hz) during rest and NREM sleep. It has been reported that hippocampal replay during SWR can be accompanied by the cortical replay of the same event (Ji and Wilson, 2007). Interestingly, optogenetic stimulation of SWRs enhances replay and results in memory improvement (Fernández-Ruiz et al., 2019). Intracranial EEG data recorded in humans has shown that SWRs in the hippocampus are nested into the troughs of the spindle oscillations (Ngo et al., 2020; Staresina et al., 2015), suggesting that spindles might facilitate the interaction, and thus communication between neural ensembles of the hippocampus and the neocortex. Notably, this memory reactivation may strengthen long-term memory representations in the neocortex such that they can support memory retrieval independent of hippocampal engagement, a process that has been proposed to serve active systems memory consolidation in sleep (Diekelmann and Born, 2010; Klinzing et al., 2019).

Converging evidence further suggests that the content of memory reactivation can be decoded from brain activity during sleep spindles (Cairney et al., 2018; Wang et al., 2019), underlining their role in reactivation-related consolidation during sleep. Indeed, targeted memory reactivation (TMR), where sensory cues related to previous learning material are used to trigger reactivation of these learning contents during sleep, is only effective if the reactivation cues are followed by a sleep spindle, demonstrating their functional role in sleep-dependent memory consolidation (Antony et al., 2018; Schönauer, 2018).

### 1.2. Sleep spindles and different types of memory

It is well-established that sleep fosters learning and retention of both declarative and procedural memories (Diekelmann and Born, 2010; Klinzing et al., 2019), and sleep spindles have been linked to these behavioral benefits. For instance, several studies showed that sleep spindles strengthen memory for a broad range of declarative tasks including wordlist and vocabulary learning (Blaskovich et al., 2017; Gais et al., 2002; Schabus et al., 2004; Schmidt et al., 2006; Studte et al., 2017, 2015; Tamminen et al., 2010), but also visual memory for pictures (Cox et al., 2012; Ward et al., 2014), and visuospatial tasks (Clemens et al., 2006; Wilhelm et al., 2011). Likewise, spindles have been shown to facilitate procedural memory, such as finger tapping, motor sequential learning, and motor adaptation (Barakat et al., 2011a; Boutin et al., 2018; Fogel and Smith, 2006; Nishida and Walker, 2007; Simor et al., 2019; Thürer et al., 2018) and mirror tracing (Holz et al., 2012; Tamaki et al., 2008; van Schalkwijk et al., 2019). However, recent findings and reviews have started to question the robustness, replicability, and generalizability of some of these findings (Ackermann et al., 2015; Cordi and Rasch, 2021). In support of this, Ackermann and colleagues (2015) observed no significant association between sleep spindles and overnight memory retention in both declarative and procedural memory measures in over 900 healthy young participants.

If spindles are a mechanism that fosters brain plasticity and if they are directly related to sleep-dependent memory reactivation, then spindle activity during sleep should be positively related to memory consolidation. In this meta-analysis, we quantified evidence for the effect of sleep spindles on memory consolidation and assessed how strongly the extant literature is subject to publication bias. We further investigated whether the consolidation of both declarative and procedural memory is equally sleep spindle-dependent. Finally, we assessed how strongly the extant literature is subject to *publication bias* and discuss to what extent this could affect the interpretation of the obtained results.

### 1.3. Spindle characteristics and their relation to memory performance

#### 1.3.1. Spindle type and topography

Studies in the 1990s hinted at the existence of two distinct sleep spindles types, observed at 10-13 Hz (slow spindles) or 13-16 Hz (fast spindles), and peaking at frontal and centroparietal cortical sites, respectively (McCormick et al., 1997; Werth et al., 1997). The idea that sleep spindles in humans are a diverse rather than prototypical phenomenon was further supported over the last few decades in studies using functional magnetic resonance imaging (fMRI; (Schabus et al., 2007), intracranial EEG (Andrillon et al., 2011; Gonzalez et al., 2022; Piantoni et al., 2017) or M/EEG source estimation techniques (Dehghani et al., 2010; Urakami, 2008). Further, a functional differentiation between slow and fast spindles has been suggested: For instance, it has been observed that both types preferentially occur at different times of the SO cycle (Mölle et al., 2011; Staresina et al., 2015), respond differentially to pharmacological interventions (Ayoub et al., 2013), and have different phenotypic and genetic profiles (Purcell et al., 2017). Spindle topography might be directly related to the spindle type that is studied (fast or slow). Additionally, it has recently been shown that the topography of spindle amplitude can also depend on the prior learning task (Petzka et al., 2022) since spindle topographies during sleep recapitulate spatial patterns of brain activation during memory encoding, a process that benefits memory consolidation. It is currently unclear whether slow and fast spindles serve distinct functional roles in memory, and how spindle topographies are related to memory consolidation. To investigate this question further, in the present meta-analysis, we assessed whether spindle type and spindle topography modulate the strength of spindle-memory association for both declarative and procedural memory.

#### 1.3.2. Spindle measures

A final important factor to consider when investigating the effect of sleep spindles on memory is the spindle measures used. Different aspects, such as count, power, or duration (see Figure 1) may reflect distinct underlying physiological processes in the brain and may be differentially associated with behavioral outcomes. Although many studies report the relationship between sleep spindles and memory, it is currently unknown if the magnitude of these associations depends on different spindle measures. The present meta-analysis will therefore assess such differences in the strength of the relationship between sleep spindles and memory consolidation across various spindle measures. (**Fig. 1)**.

**Fig. 1.**
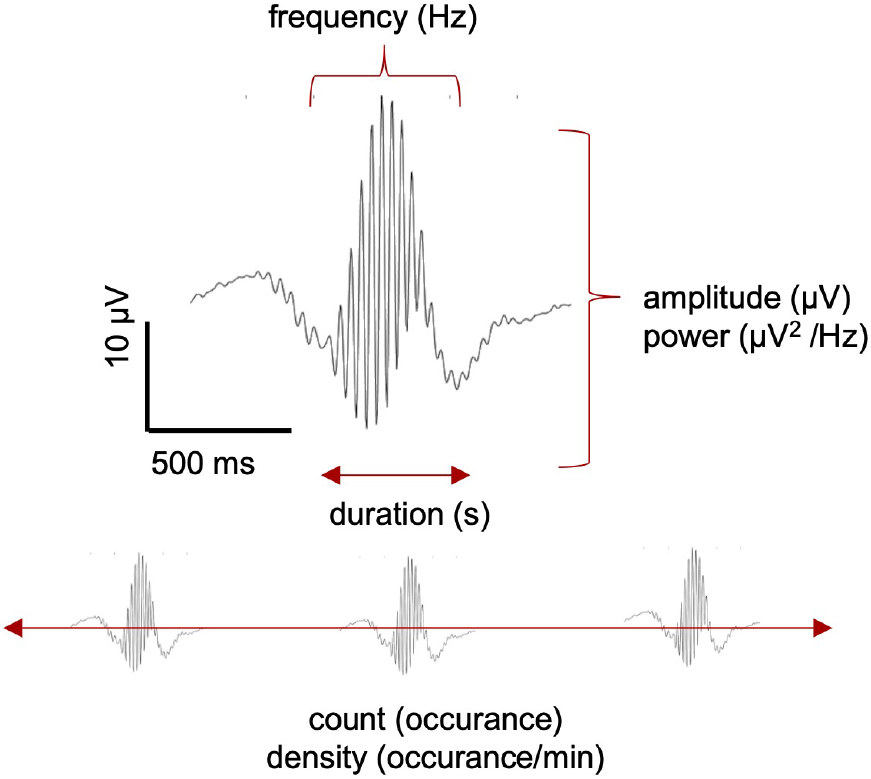
Schematic representation of how spindles appear in a non-rapid eye movement (NREM) sleep in electroencephalogram (EEG) and overview of the most frequently studied measures. *Count*: number of spindles; *Duration:* mean time (in sec) of spindle events; *Density*: number of spindles per unit of time (per minute or per 30 sec); *Power*: mean amount of activity in spindle frequency; *Amplitude:* the absolute value (in μV) of the maximum negative peak, measured as the voltage difference between the largest trough and the largest peak; *Frequency*: Number of occurrences of a repeating event per unit of time (cycle number/duration) or maximal spectral peak amplitude (in Hz); *Spindle Activity (SpA):* mean duration x amplitude.

### 1.4. Objectives of the meta-analysis

Although there are meta-analyses investigating the relation of sleep spindles with trait-like aspects like intelligence, cognitive performance, and general cognitive ability in healthy adults (e.g. Ujma, 2018), in the present study, we aim to contribute a more comprehensive overview of the relationship between sleep spindles and memory. We review the relevant methodological aspects that vary across studies and report meta-analytical evidence for the relationship between sleep spindles measured with EEG and memory. More precisely, we aimed to address the following main research questions systematically:

1. What is the overall strength of the association between sleep spindles and memory performance?
2. Does the sleep spindle-memory association depend on the type of memory that is consolidated (e.g., procedural vs. declarative)?
3. Do certain spindle types and spindle scalp locations result in stronger spindle-memory associations?
4. Does the strength of the sleep spindle-memory relationship depend on the spindle measures?

To investigate the general effect of sleep spindles on memory, we included studies reporting different kinds of memory measures. Both memory retention and post-sleep memory measures were categorized as reflecting the process of memory consolidation, and thus constitute a *state* measure of memory influenced by the specific learning experience. *Trait-like* measures of memory, such as recall performance on a control day or pre-sleep memory performance were categorized as learning ability, reflecting a general cognitive trait that is largely independent of the specific learning experience (Clemens et al., 2006, 2005; Lustenberger et al., 2012; Schabus et al., 2008; Sopp et al., 2018). In the main part of the meta-analysis, we focused on the association between sleep spindles and memory consolidation as a *state* measure and assessed how various spindle characteristics influence this relationship, but we also evaluated the overall strength of the spindle-memory association for the *trait* measure. We finally explored the strength of the spindle-memory association depending on the sleep protocol and sleep duration, as the included studies investigated sleep both during the whole night and nap sleep protocols.

## 2. Methods

### 2.1. Search strategy

The present meta-analysis was implemented according to the Preferred Reporting Items for Systematic Reviews and Meta-Analyses (PRISMA) guidelines for designing, implementing, and reporting systematic reviews and meta-analyses (Page et al., 2021). The systematic literature search was conducted on electronic databases including PubMed and PsychINFO in May 2021 using the following terms: sleep spindles, memory, EEG as well as database-specific search terms (including Medical Subject Heading (MeSH) terms). Additionally, further sources including reference lists of included studies and previous systematic reviews on the study topic as well as two key journals (Neuroscience & Biobehavioral Reviews and Sleep Medicine Reviews) were searched to increase the likelihood of retrieving relevant empirical studies.

### 2.2. Study selection

Studies were included if they **1)** reported results from a cohort, case-control, cross-sectional or experimental study and **2)** assessed the relationship between memory (procedural and/or declarative) and sleep spindles measured with EEG in healthy individuals between 18 and 65 years of age. Additionally, studies met the inclusion criteria only if they were published in peer-reviewed journals in English. On top of the inclusion criteria, we excluded studies with **1)** non-human samples (e.g., rodents, primates), **2)** samples from clinical populations (e.g., neurological or psychiatric disorders), **3)** the use of a reactivation protocol including targeted memory reactivation (TMR), **4)** applying pharmacological (e.g., SSRIs) or physiological interventions (e.g., transcranial direct current stimulation (tDCS), transcranial magnetic stimulation (TMS), auditory simulations), **5)** the absence of a long-term memory measure (e.g., reporting only working memory or measures of executive function), or **6)** absence of sleep, EEG, or spindle measures. Other types of studies including systematic reviews, meta-analyses, letters to editors, opinions, case reports, case series, and clinical trials were also excluded.

### 2.3. Study screening and data extraction

Identified titles and abstracts were screened for relevance, separately by the two reviewers (DK, AM). Full texts of identified articles were retrieved and read in full to assess eligibility for inclusion. A standardized data coding form was developed to extract the following information from each study: **(a)** authors, publication year, and DOI; **(b)** characteristics of the study sample (age, gender, number of participants); **(c)** spindle characteristics (type and measure); **(d)** sleep information (duration, protocol, stage); **(e)** EEG information (scalp location, channel); **(f)** memory type and specific outcome measure and **(g)** statistics (e.g., the correlation between memory and sleep spindles).

In cases of relevant studies not reporting extractable statistics with regard to the association between sleep spindles and memory performance, authors were contacted via email to request the minimum required data for the meta-analysis. Emails were sent at least twice at approximately fourteen-day intervals.

Importantly, we aim to make this meta-analysis transparent by publicly sharing data files, associated codebooks, and analysis at Open Science Framework https://osf.io/wu6d7.

### 2.4. Statistical Analysis

The meta-analysis was conducted using the *metafor* package (Viechtbauer, 2010) with the function of *rma.mv* in the *R* statistical software (version 3.5; R Core Team, 2017), allowing to fit multilevel meta-analytic models.

Before computing the meta-analyses, correlation coefficients were transformed to the normally distributed Fisher’s z (*escalc*). The Fisher’s z score and its variance were used in the meta-analyses, and afterwards back-transformed to Pearson’s correlation coefficients to facilitate interpretation. If Pearson correlations were not reported, Fisher’s z was estimated using statistics such as partial r and Spearman’s rank correlation.

Since the studies included in this meta-analysis have dependent effect sizes, with several effect sizes reported from the same sample, statistical analysis based on traditional approaches such as averaging all effect sizes, selecting one effect size per study, or conducting separate meta-analyses in independent subsets may lead to inaccurate or false-positive results (Borenstein et al., 2009; Cheung, 2019). The extant literature has proposed several procedures to account for these dependencies (Borenstein et al., 2009). In the current study, we opted for a multilevel random-effects model using a restricted maximum likelihood method (REML) to estimate overall effects and account for the heterogeneity across included studies (Cheung, 2019; Fernández-Castilla et al., 2020; Konstantopoulos, 2011). The three-level randomeffects model quantifies the sampling variance (level 1), the variance between effect sizes within studies (level 2), and the variance of effect sizes between studies (level 3). To examine variance and heterogeneity among effect sizes of included studies, we computed *Q* statistics, the significance of *Q*, and *I*^2^, respectively. *I*^2^ values of ~25%, 50%, and 75% were interpreted as low, moderate, and high, respectively (Higgins and Thompson, 2002; Quintana, 2015; Thompson and Higgins, 2002). Lastly, the presence of publication bias was examined by investigating the association between sleep spindles and memory consolidation with the modified version of the Egger test using standard error as a moderator. To estimate how publication bias may influence the estimated effect sizes between sleep spindles and memory, we used information on whether an effect size was originally reported in the literature or whether it was obtained only later, upon asking the authors (effect size reporting status: reported vs. unreported) as a categorical moderator to assess whether reported correlation values had significantly larger effect sizes than unreported ones (for a similar approach, see (Hu et al., 2020)).

### 2.5. Moderators

A first moderator analysis was conducted to determine the overall strength of the association between sleep spindles and memory performance depending on state consolidation and trait learning ability measures. With regard to memory consolidation measures, we then assessed possible effect size differences between memory retention and post-sleep memory measures. We further investigated differences in the magnitude of sleep-memory associations for declarative compared with procedural memory. As sleep may contribute differentially to the memory consolidation depending on the procedural memory task (Schendan et al., 2003; Schönauer et al., 2015; Walker et al., 2003), we also explored effect size differences between hippocampal (e.g., finger tapping) vs. non-hippocampal procedural tasks (e.g., mirror-tracing or motor-adaptation). We next assessed the effect of spindle type and EEG scalp topography on memory consolidation. Spindle type was defined as slow, fast, and global spindle-frequency range. For investigating scalp topography, we grouped EEG channels into three coarser brain regions (frontal, central, and parietal), as the EEG montage system, electrode positioning, and the number of electrodes varied between studies. We additionally explored the effect of EEG scalp topography depending on spindle type, as it was previously demonstrated that slow spindles prevail over anterior and fast spindles over posterior brain areas (Cox et al., 2017). We finally examined differences in the spindle-memory association depending on spindle measures, which were categorized as power, duration, count, amplitude, density, oscillatory frequency, and spindle activity (SpA) (**Fig. 1)**. We defined these variables based on the definition or calculation reported in the original studies. In addition to these planned analyses, we also explored the possible effects of sleep protocol (whole night vs. napping), sleep duration (in min), and a possible effect of age (in years), as these variables may impact sleep-behavior correlates (Gui et al., 2017; Hokett et al., 2021; Lo et al., 2016; Qin et al., 2022; Schmid et al., 2020). In all analyses, moderator effects for categorical variables were only considered if at least six to seven effects sizes were available per category (Tipton et al., 2019). In all analyses, moderator effects for categorical variables were only considered if at least six to seven effects sizes were available per category (Tipton et al., 2019).

### 2.6. Control Analyses

Two control analyses were conducted to compare meta-analytic results obtained from analyses with and without outliers, as the presence of outliers may affect the validity and robustness of the conclusions from a meta-analysis (Viechtbauer and Cheung, 2010). We determined statistical outliers using standardized residuals and Cook’s distances (*D_i_* (Cook, 1977). *D_i_* indicates the relative influence of each effect size on the summary estimate. A standard rule of thumb for potential outliers is a Di value greater than three times the mean *D_i_*(Viechtbauer and Cheung, 2010). Similarly, as an outlier criterion, we excluded effect sizes that have standardized residual greater than 3 in absolute magnitude (Hedges and Olkin, 2014).

As a second control analysis, we excluded studies with the largest sample size (Ackermann et al., 2015) and the smallest sample size (Fogel and Smith, 2006) and also on studies that reported a substantially larger number of estimates than the average (Blaskovich et al., 2017; Thürer et al., 2018; k > 130) and repeated the meta-analysis for the association between sleep spindles and memory consolidation.

## 3. Results

### 3.1. Characteristics of the included studies

After reviewing the 421 titles and abstracts, 140 studies were retained for full-text review and screened for eligibility. Eighty-seven of these studies were excluded for the following reasons: Not providing a spindle characteristics or type diagnosis (n = 12), no clear measure of memory (n = 35), no sleep (n = 2) or EEG acquisition (n = 4), lacking necessary data to compute effect sizes (n = 25), investigating dream-recall (n = 2), working memory/intelligence (n = 6) or other (n = 1) as shown in **Fig. 2.** A total of 53 studies were included in the final analysis published between 2002 (Gais et al., 2002) and 2021 (Lutz et al., 2021) (**Fig. 3)**. The included studies were conducted in the following countries: Germany (n = 14), United States of America (n = 8), Canada (n = 7), Austria (n = 6), Switzerland (n = 4), United Kingdom (n = 4), Hungary (n = 4), The Netherlands (n = 2), Japan (n = 2), Iran (n = 1), and Cuba (n = 1). There were 33 studies investigating declarative memory, 16 studies assessing procedural memory, and 4 studies reporting results from both (Ackermann et al., 2015; Genzel et al., 2014; Griessenberger et al., 2013; Holz et al., 2012). Further, there were 6 putatively non-hippocampal procedural tasks like mirror-tracing (e.g., Tamaki et al., 2008; van Schalkwijk et al., 2020) and 14 procedural tasks that have been shown to rely on hippocampal contributions, like finger sequence tapping (e.g., Nishida et al., 2016; Schönauer et al., 2014).

**Fig. 2.**
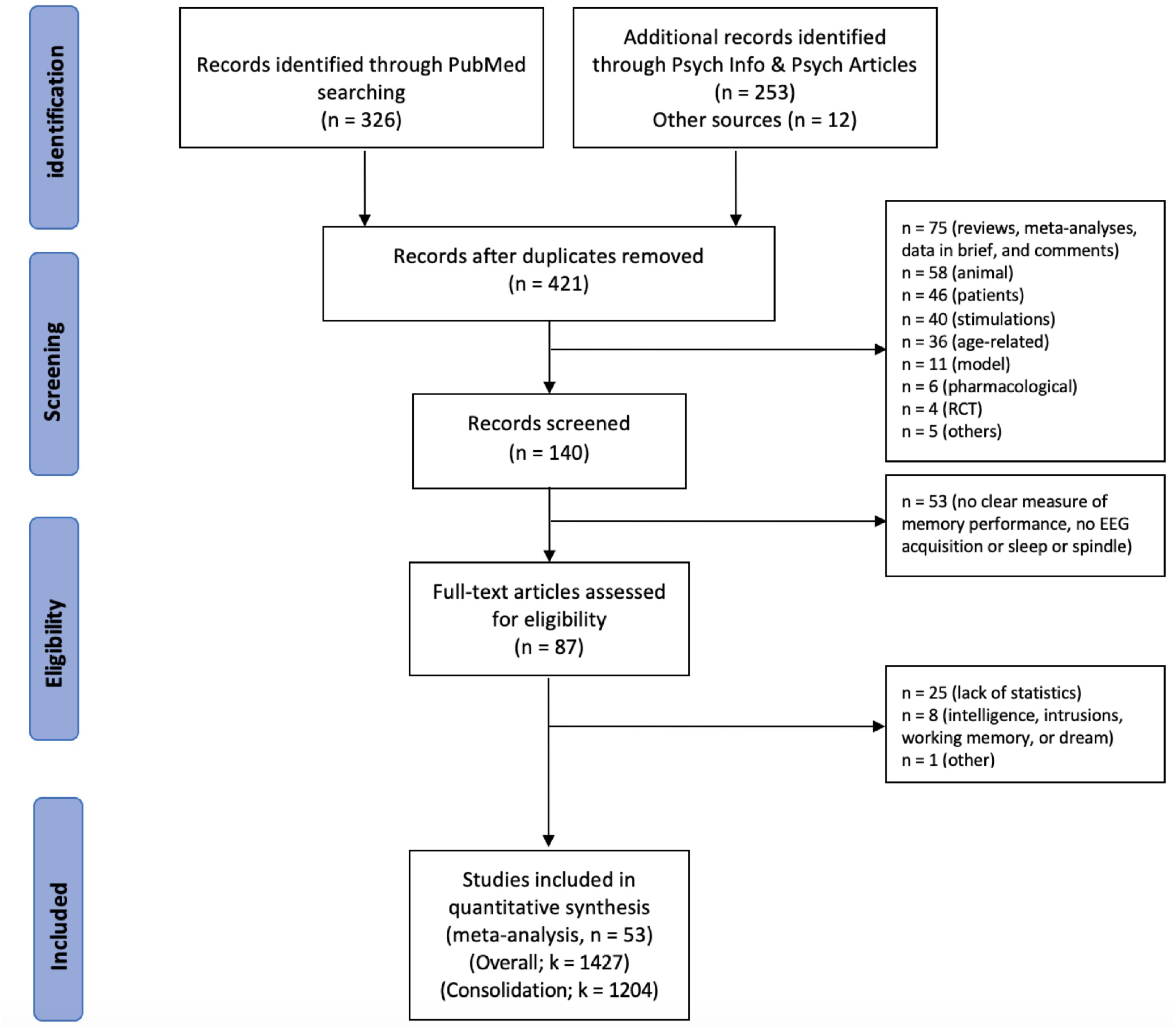
PRISMA flow diagram of the literature search, screening, and inclusion processes; n = number of studies; k = number of estimates.

**Fig. 3.**
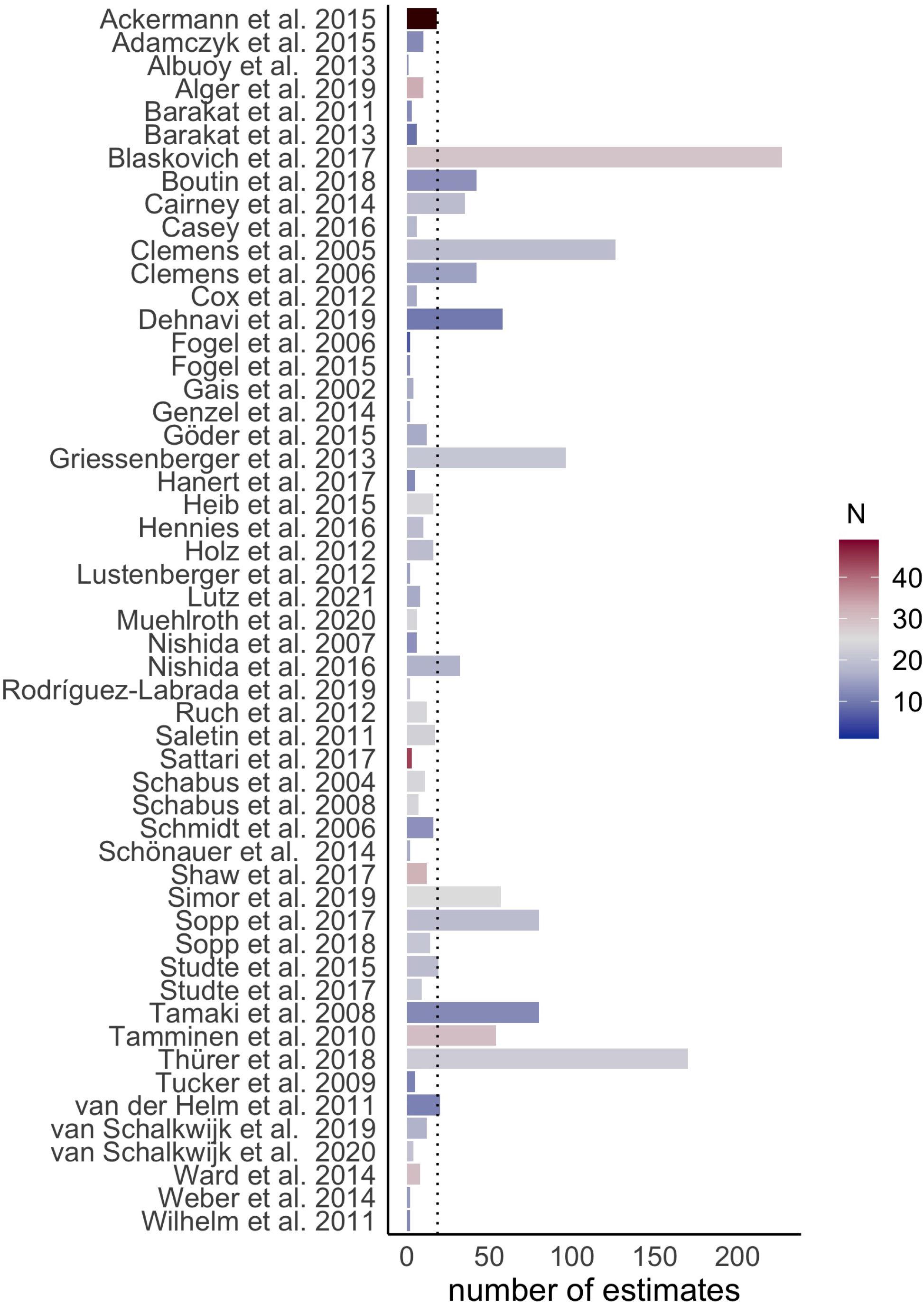
Overall number of estimates and sample size per study extracted for the meta-analyses (k=1427). To increase the interpretability of the graph, we visualized one study (Ackermann et al., 2015) with the largest sample size (N = 929) as dark brown. While the average sample size (N) across 53 studies was 35.66, after excluding this study the average sample size became 18.59 (ranging from N = 6 to N = 44), shown in vertical dot line. The total sample size across 53 studies was 1896.

The number of estimates for each measure of interest across 53 studies in 1427 effect sizes can be found in Supplementary Table 1.

### 3.2. The association between sleep spindles and memory performance

We first tested the overall strength of association between sleep spindles and memory performance including both state consolidation and trait learning ability measures in 1427 unique associations (k) across 53 independent studies. Using a three-level meta-analytic model, we observed that the pooled correlation between sleep spindles and memory performance was r = 0.236; CI: [0.161–0.308]; *p* < 0.001, reflecting a small- to moderate effect size for the spindle-memory association. Moderation analyses indicated that there were no differences in effect sizes of the spindle-memory association between state and trait measures of memory (*Q_m_*(1) = 0.560, *p* = 0.454), but we observed a numerically higher effect size for state consolidation measures (r = 0.238, k = 1204) than for trait-like measures (r = 0.218, k = 223).

### 3.3. Average association between sleep spindles and memory consolidation

Since our main objective was to assess spindle-dependent memory consolidation and how various spindle characteristics, including spindle type, spindle topography, and different spindle measures, influence the spindle-memory relationship, we focused on state consolidation measures from this point onwards.

We first tested the strength of association between sleep spindles and memory consolidation and observed a moderate correlation (r = 0.243; CI: [0.166–0.317]; *p* < .001; Supplementary Fig 1). There was substantial heterogeneity among effect sizes (*Q*(1203) = 2828.262, *p* < .001; *I*^2^(2) = 21.43 and *I*^2^(3) = 52.75 especially moderate to high between-study heterogeneity. We assessed publication bias using the modified version of the Egger test using standard error as moderator and detected significant publication bias (β = 1.64, *SE* = 0.53, *p* = 0.0020, Supplementary Fig 2, Supplementary Fig 3), indicating that significant results are more likely to be published than small or non-significant findings. We then tested whether the effect sizes were larger if reported in published manuscripts than if they were not reported. When effect size reporting status (yes vs. no) was examined in the moderator analysis, we observed no significant differences between reported and unreported effect sizes (*Q_m_*(1) = 0.445, *p* = 0.505; reported; r = 0.249, k = 793, unreported; r = 0.222, k = 441) across all studies. Interestingly, when we conducted this moderation analyses separately for studies assessing declarative or procedural memory, we observed significant differences (*Q_m_*(1) = 6.094, *p* = 0.014 in declarative; (*Q_m_*(1) = 13.645, *p* < 0.01 in procedural memory): While in studies with declarative memory we found the effect size of reported studies (r = 0.254, k = 517) were higher than unreported ones (r = 0.131, k = 295), we found opposite pattern such that the effect size of reported results (r = 0.169, k = 276) were higher than unreported ones (r = 0.421, k = 116) in studies with assessing procedural memory.

Subsequent moderation analyses further revealed no differences between memory retention and post-sleep measures (*Q_m_* (1) = 0.341, *p* = 0.560; memory retention; r = 0.249, k = 592, post-sleep; r = 0.231, k = 612), suggesting that both measures show a similar effect size for the spindle-memory association.

### 3.4. Spindle-memory association depending on memory type

To address the second research question, memory type (declarative vs. procedural) was examined as a moderator for the mean effect size between sleep spindles and memory consolidation. We found that memory type is a significant moderator of the spindle-memory association (*Q_m_*(1) = 7.589, *p* = 0.006), with a stronger spindle-memory association for procedural memory (r = 0.319, *p* < .001, k = 392) than for declarative memory (r = 0.210, *p* < .001, k = 812), as shown in (**Fig. 4A**. Although studies employing hippocampal procedural memory tasks (r = 0.263, k = 195) showed higher effect sizes than those using non-hippocampal tasks (r = 0.244, k = 197), this was not a significant moderator (*Q_m_*(1) = 0.008, *p* = 0.928).

**Fig. 4.**
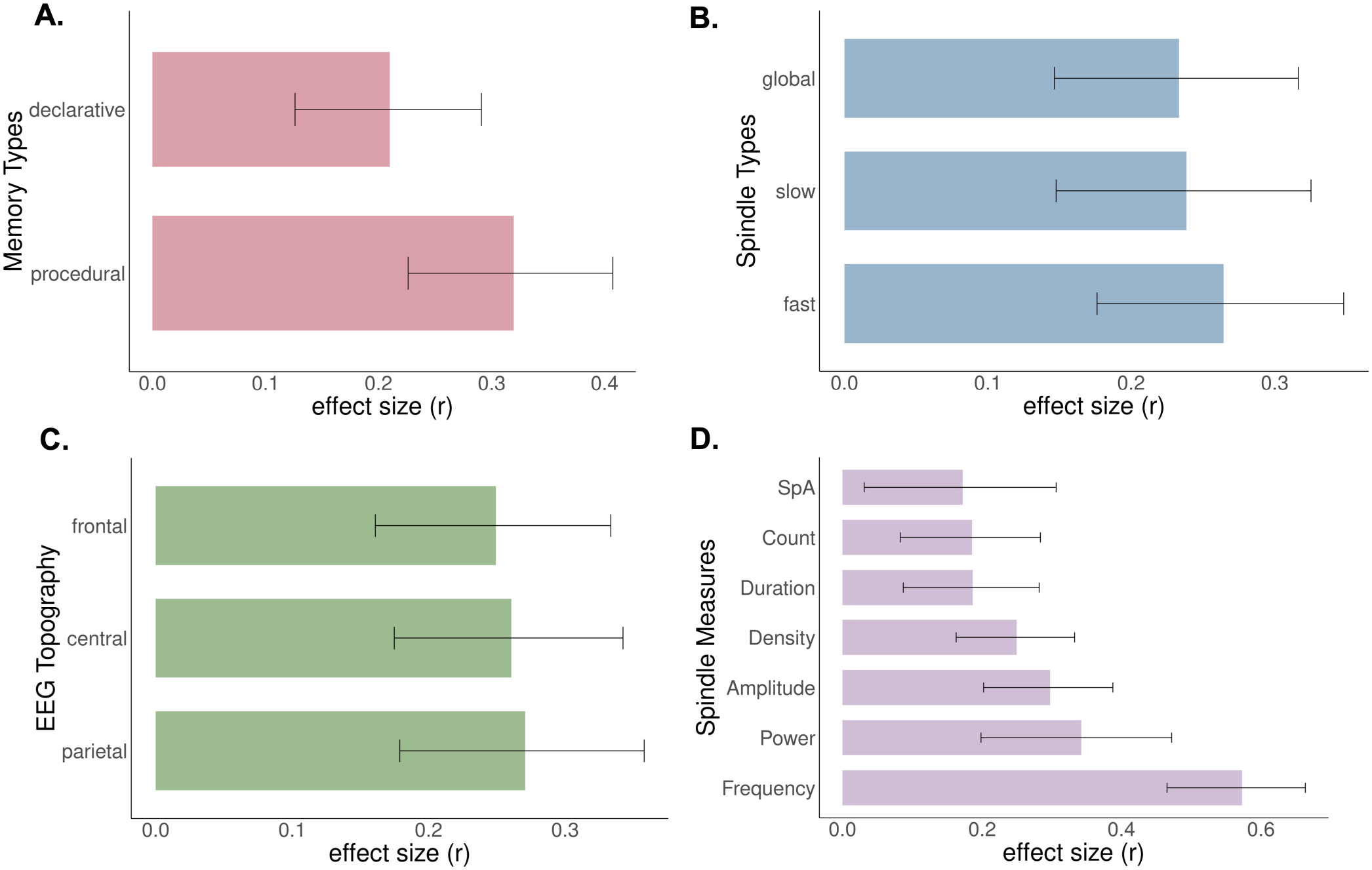
**The average effect size (r) of sleep-memory associations by A**. memory types, **B**. spindle types, **C**. EEG topography, and **D**. spindle measures. Error bars indicate 95% confidence intervals. While we observed significant effect size differences for memory types and spindle measures, there were no significant moderation of spindle type and scalp topography on spindle-memory consolidation.

### 3.5. Spindle-memory association depending on spindle characteristics

#### 3.5.1. Spindle types

We further assessed whether spindle type, i.e., slow or fast spindles, acted as a moderator for spindle-memory associations across all studies. There was a no moderation effect of spindle type (*Q_m_*(2) = 1.831, *p* = 0.400; fast; r = 0.264, k = 552, slow; r = 0.238, k = 327, global; r = 0.233, k = 325, **Fig. 4B**). The subsequent moderation analysis separately performed in studies on declarative memory was also non-significant (*Q_m_*(2) = 0.675, *p* = 0.714, k = 812), suggesting no notable difference between the spindle types in declarative memory (fast; r = 0.244, k = 308, slow; r = 0.250, k = 226, global; r = 0.216, k = 278). Nevertheless, we found a significant effect of spindle type (*Q_m_*(2) = 8.610, *p* = 0.013, k = 392) for procedural memory measures, where global and fast, but not slow frequency spindles contributed to the spindle-memory relationship (fast; r = 0.247, *p* = 0.01, k = 244, slow; r = 0.141, *p* = 0.186, k = 101, global; r = 0.328, *p* = 0.005, k = 47). Follow up post-hoc comparisons with Bonferroni correction indicated a correlation difference between fast and slow (fast-slow; rdiff = 0.110, (*p_adj_* = 0.007), but not between fast and global or slow and global spindle frequencies in procedural memory (*p_adj_* > 0.05).

#### 3.5.2. EEG scalp topography

Next, moderator analyses were conducted to determine if scalp topography (frontal (k = 350), central (k = 438), or parietal (k = 222) modulated the strength of the spindle-memory relation. There was no significant moderation effect of scalp topography across all studies (*Q_m_*(2) = 0.885, *p* = 0.659, k = 1010), meaning that there was no influence of recording location of the spindles on the strength of the spindle-memory associations (**Fig. 4C**). Moreover, we calculated separate metaanalyses across studies reporting slow and fast spindles, respectively and observed no moderator effect of scalp topography (slow; (*Q_m_*(2) = 0.186, *p* = 0.911, k = 297, fast; (*Q_m_*(2) = 0.137, *p* = 0.934, k = 488), suggesting that the effect sizes of EEG clusters do not differ in studies reporting either slow or fast spindle frequencies (r = 0.125 – 0.175).

#### 3.5.3. Spindle measures

We further investigated effect size differences in the sleep-memory association depending on different spindle measures (amplitude; k = 142, count; k = 235, density; k = 321, duration; k =145, frequency; k = 27, power; k = 204, and SpA; k = 130). We found a significant moderation effect of spindle measure (*Q_m_*(6) = 69.692, *p* < 0.001, k = 1204) with the largest effect sizes for frequency and power (**Fig. 4D**). All of the spindle measures showed significant memory associations (all *p* < 0.05). The results of the post-hoc tests (number of comparisons = 21) indicated larger effects for spindle frequency compared to all spindle measures except for spindle power (*p* = 0.06) and amplitude compared to duration (*p_adj_* < 0.05, Supplementary Table 2), while we did not observe any other significant effect size differences between the rest of the spindle measures after Bonferroni correction (all *p_adj_* >0.05, Supplementary Table 2).

Because only two studies reported the relation of memory consolidation (procedural memory) with oscillatory peak frequency (Boutin et al., 2018; Thürer et al., 2018), defined as maximal spectral peak amplitude (in Hz), we repeated the analysis after excluding the frequency measure from these studies (n = 27). After this exclusion, we confirmed the observed findings of the moderator effect of spindle measures (*Q_m_*(5) = 14.968, *p* = 0.01, k = 1177) with the largest effect size for the spindle power and the smallest effect size for SpA. Further post-hoc comparisons (number of comparisons = 15) similarly indicated that there were significant effect size differences between amplitude and duration, but not between other measures (*p_adj_* < 0.05, Supplementary Table 3).

### 3.6 Exploratory Analyses

#### 3.6.1. Sleep Protocol and Duration

Moderator analyses were conducted to determine if sleep protocol (nap vs. whole-night) or sleep duration (as a continuous variable) modulated the strength of the sleep-memory association. There was no significant difference (*Q_m_*(1) = 2.149, *p* = 0.143, k = 1169) in sleep-memory associations for nap sleep (r = 0.167, k = 543) and whole night sleep (r = 0.280, k = 626). Confirming this result, we also observed non-significant findings in studies reporting either declarative (*Q_m_*(1) = 0.471, *p* = 0.493, k = 812) or procedural memory measures (*Q_m_*(1) = 2.089, *p* = 0.148, k = 357). Similarly, we found no significant moderation effect of sleep duration (M = 242.41, SD =169 min) on spindle-memory association neither across all studies (*Q_m_*(1) = 1.357, *p* = 0.244, k = 1204, r = 0.181), nor separately in studies reporting either declarative (*Q_m_*(1) = 1.032, *p* = 0.310, k = 812, r = 0.174) or procedural memory (*Q_m_*(1) = 0.251, *p* = 0.617, k = 392, r = 0.210).

#### 3.6.2. Age

Moderator analyses were conducted to determine if age (as a continuous variable) modulated the strength of the sleep-memory association. We observed that continuous age (in years) was not a significant moderator for the association between sleep spindles and memory (*Q_m_*(1) = 0.235, *p* = 0.628, r = 0.355). We observed similar results when separately analyzing studies reporting declarative (*Q_m_*(1) = 0.204, *p* = 0.652, r = 0.138) and procedural memory (*Q_m_*(1) = 1.100, *p* = 0.294, r = 0.744).

### 3.7. Control Analyses

After computing standardized residuals and Cook’s distance (Di), 50 effect sizes were excluded. We observed similar findings for the summary estimate of the spindle-dependent memory consolidation and no major changes occurred to the results of moderator analyses.

After excluding studies with the largest (Ackermann et al., 2015) and smallest (Fogel and Smith, 2006) sample size and a higher number of estimates (Blaskovich et al., 2017; Thürer et al., 2018), no substantial changes occurred to the main summary statistics with memory consolidation measures. However, we observed a major shift in the moderation results. Specifically, the effect of scalp topography was statistically significant in these analyses, whereas we no longer found a significant effect of spindle measure on memory consolidation. Given that we removed one-third of the estimates and sample size, such statistical differences are to be expected. The details of both analyses can be found online (https://osf.io/wu6d7). In this article, we focus on the initial findings for the spindle-dependent memory consolidation using all effect sizes (k = 1204).

## 4. Discussion

The present meta-analysis aimed to **i)** determine if there is sufficient evidence to support the claim that sleep spindles facilitate memory processing and **ii)** assess how multiple factors, including memory type, spindle characteristics, and EEG scalp topography affect the sleep spindle-memory association. The literature search yielded 53 studies that were quantitatively analyzed. We found a small to moderate effect size for the association between sleep spindle and memory measures. Moderation analyses revealed that the association between sleep spindles and memory consolidation was stronger for procedural memory than for declarative memory. Neither spindle type (i.e. fast vs. slow) nor EEG scalp topography impacted the strength of the spindle-memory association, but we found a moderator effect of spindle measures, with the largest effect sizes for spindle frequency and spindle power. Exploratory analyses indicated no differences in effect size for the spindle-memory association between sleep protocols (nap vs. whole-night sleep) or sleep duration, indicating that sleep spindles may be a general mechanism supporting memory consolidation during sleep.

### 4.1. Sleep spindles and memory consolidation

The studies included in this meta-analysis reported both associations of trait learning abilities with spindle measures, and how sleep spindles are related to state-dependent consolidation of newly learnt material. We found that sleep spindles show a consistent relationship with both trait-like and state-like measures of memory, with a numerically higher effect size for state-dependent consolidation measures. We focused our analyses on a systematic investigation of sleep spindles on the consolidation of newly learnt information. Theoretical accounts of sleep-dependent memory consolidation postulate an important role of sleep spindles in memory consolidation. Mechanistically, the consolidation of declarative memories in NREM sleep is thought to depend on the interaction of characteristic oscillatory signatures, including sleep spindles. As proposed in the model of active systems consolidation during sleep, brain-wide reactivation and integration of hippocampal memories requires global synchronization and communication between cortical and subcortical neural networks (Diekelmann and Born, 2010; Klinzing et al., 2019; Maingret et al., 2016). Sleep spindles, which are generated in thalamo-cortical circuits, may facilitate this dialogue. Cortical neurons are excited by thalamocortical neurons which generate spindle activity in the cortical field potential, while the neocortex in turn drives spindle onset and regulates their termination via projections to the thalamus (Timofeev et al., 2001; Timofeev and Chauvette, 2013), indicating a coordinated communication between cortical and subcortical structures. This dialogue between the thalamus and neocortex is instrumental to memory functions: optogenetic stimulation of thalamic sleep spindles drives cross-regional spindle co-occurrence and supports hippocampusdependent learning (Latchoumane et al., 2017). Sleep spindles may thus be actively involved in the dialogue between segregated brain areas in the service of memory consolidation. Our finding that sleep spindles support memory performance, and the consolidation of newly learnt memories in particular, is thus in line with theoretical accounts of sleep-dependent memory consolidation and findings that implicate spindles in nighttime reactivation of memories. Although we could not directly determine the neural mechanisms by which sleep spindles exert their influence on memory functions, we systematically investigated potential electrophysiological and behavioral moderators to further characterize the spindle-memory relationship.

### 4.2. Spindle-dependent memory consolidation in different types of memory

In the current study, we observed that EEG sleep spindles are associated with memory consolidation in both major memory domains, consistent with the idea that sleep spindles are not only implicated in declarative memory (Peyrache and Seibt, 2020; Schabus et al., 2004) but also in procedural memory and skill learning (Boutin and Doyon, 2020; King et al., 2017; Walker et al., 2005). Interestingly, the strength of spindle-memory association was stronger for procedural than for declarative memory. Previous research on sequential motor learning tasks demonstrated that increases in BOLD signal in the cortico-striatal system during learning were correlated positively with both fast and slow spindle activity during later sleep (Barakat et al., 2013). As these tasks rely on hippocampal activity, next to the striatum, precuneus, prefrontal cortex, and primary motor cortex (Albouy et al., 2015, 2008; Boutin et al., 2018; King et al., 2017), a network of brain regions that overlaps with areas involved in declarative memory, the role of sleep spindles for the consolidation of sequential motor learning may be functionally similar to consolidation in the declarative memory system. Indeed, systems memory consolidation has recently been observed in the procedural domain (Boutin and Doyon, 2020; King et al., 2017; Schmid et al., 2020). Importantly, sleep spindles have been shown to foster changes in the neural substrate supporting memories and to promote brain-wide integration after learning in both declarative (Bergmann et al., 2012; Cowan et al., 2020; Jegou et al., 2019) and procedural memory tasks (Boutin et al., 2018; Fogel et al., 2017). While historically the role of spindles has mainly been discussed for declarative memory, future theoretical accounts should thus consider that similar mechanisms may apply in the procedural domain.

Because hippocampal coordination is a central factor in theoretical accounts of sleep-dependent consolidation in the declarative domain, yet many procedural tasks do not require hippocampal activity during acquisition, we further tested whether sleep spindles equally contribute to procedural memory tasks with and without hippocampal involvement (Schendan et al., 2003; Walker et al., 2003). We observed no moderator effect of procedural memory task (non-hippocampal vs. hippocampal) on spindle-memory associations. This is in line with a recent meta-analysis that similarly reported a small effect of sleep on motor memory consolidation with no notable differences between various motor tasks including finger tapping and mirror tracing (Schmid et al., 2020). This is surprising, given an elaborate theoretical and experimental foundation that links spindle-related consolidation to a cross-regional dialogue between the hippocampus, thalamus, and neocortex (Antony et al., 2019; Latchoumane et al., 2017; Staresina et al., 2015) and stresses the role of the hippocampus reactivation-dependent consolidation. It has been suggested that the hippocampus could play a broader role in tasks traditionally considered hippocampal-independent, where it may help to reinstate extra-hippocampal memory traces (Sawangjit et al., 2018). Therefore, both hippocampal and non-hippocampal procedural memory tasks may indeed benefit similarly from spindle activity during sleep. Alternatively, sleep spindles may benefit memory via mechanisms independent of hippocampally coordinated reinstatement of learning-related activity. Indeed, it has been found that the thalamus can instruct plasticity in the primary visual cortex during a sleep period following visual discrimination learning and that sleep spindles mediate this effect (Durkin et al., 2017). Spindles’ role in memory may thus be two-fold: In supporting thalamo-cortical and thalamo-hippocampal communication, they may aid systems-level consolidation, but their plasticity-promoting properties can also foster synaptic consolidation, more generally. Because also memory tasks with no hippocampal contribution showed a spindle-related consolidation benefit, we would argue that our results support this more global role for spindles in mnemonic processes such as consolidation.

### 4.3. Spindle characteristics and their relation to memory consolidation

#### 4.3.1. Spindle Types and Topography

As mentioned in the introduction, many authors segregate sleep spindles into two main types according to their oscillatory frequency and topography, with slow spindles occurring over frontal regions and fast spindles dominating over parietal and central sites (Andrillon et al., 2011; Cox et al., 2017; Schabus et al., 2007; Urakami, 2008). It has further been hypothesized that slow frontal spindles may be preferentially generated by cortico-cortical or intracortical mechanisms, whereas faster parietal spindle activity is more closely related to thalamo-cortical communication (Ayoub et al., 2013; Timofeev and Chauvette, 2013). However, the distinct functional roles of these spindle types remain largely unknown. Given that spindle oscillations are widely distributed across the brain, it may seem plausible to assume that sleep spindles foster the strengthening of memory traces in different memory systems depending on spindle type (fast vs. slow), their location, or both. In the present meta-analysis, we assessed this question by examining the moderator effect of spindle types and of EEG scalp topography. Although spindle type and scalp topography did not impact the strength of the overall spindle-memory association, we observed that the procedural memory-spindle association was moderated by spindle types, with global and fast sleep spindles, showing the strongest effect. This result is in line with previous studies, which demonstrated that global and fast sleep spindles are related to memory improvement in motor procedural memory tasks (Barakat et al., 2013, 2011b; Fogel and Smith, 2006; Nishida and Walker, 2007; Simor et al., 2019; Tamaki et al., 2008). As fast sleep spindles dominate over parietal and central sites including motor regions, we believe that our results may reflect learning-related patterns of brain activation underlying localized spindle expression during sleep (for a recent study demonstrating this in declarative learning, see Petzka et al., 2022).

Importantly, in the literature, there is no clear consensus on the spectral distinction between fast and slow sleep spindles (De Gennaro and Ferrara, 2003). While most studies have highlighted a distinct scalp topography for fast and slow spindles (Cox et al., 2017; Schabus et al., 2007), several experiments reported overlapping M/EEG sources for faster and slower spindle components (Dehghani et al., 2010; Gumenyuk et al., 2009). Moreover, unlike in humans, a distinction between slow frontal and fast parietal spindles in mean frequency is not found in mice (Kim et al., 2015).

The way in which spindles manifest on the scalp might be linked to the structural anatomy of the generating network (Piantoni et al., 2016). In other words, the frequency variability might be caused by differences in anatomical structure, like the length or degree of myelination of thalamo-cortical white-matter tracts (Piantoni et al., 2017, 2013), which affect conduction velocity (Fields, 2015) and axonal propagation delays (O’Reilly and Nielsen, 2014). Alternatively, task-specific localization of neuronal activity may drive functional differences between spindle subtypes and differences in spindle topography. Speaking against the idea that slow and fast spindles rely on a distinct neural circuitry, a recent electro-corticography study with laminar profiling in humans suggested that slow and fast spindle subtypes are generated in cortical and thalamic circuits that have similar projections to cortical regions (Ujma et al., 2021). Although in the present meta-analysis, we did not observe effect size differences depending on EEG scalp location, our results do not contradict the notion that the topography of sleep spindles can reflect or even bear relevance to memory consolidation. A recent study suggests that participant-specific topographies of sleep spindles are related to regional activation during memory encoding, and spindle activity during later sleep may be relevant for expressing plasticity in these regions (Petzka et al., 2022). Our topographical analyses do not reflect such subtle spatial patterns. Given what we know about spindle generation and their role in plasticity, determining their regional expression, however, appears a promising area for future research.

#### 4.3.2. Spindle Measures

Spindle-memory associations can be measured using various sleep spindles properties including amplitude, count, density, duration, frequency, power, and SpA. However, it is still unknown whether the magnitude depends on specific spindle measures, which may reflect both similar or different biological underpinnings. In the present meta-analysis, we assessed a moderation effect of spindle measure and expected that the spindle-memory association would show a similar strength for different spindle measures, given their high intercorrelation (O’Reilly and Nielsen, 2014). Yet, we observed that spindle measures moderated the spindle-memory association with the strongest effect for spindle frequency and spindle power.

Our research found a strong effect of spindle peak frequency on memory consolidation. However, it is important to note that this finding should be interpreted with caution as there have been only two studies (k=27) reporting a correlation between spindle peak frequency and memory consolidation. Individual spindle peak frequency is known to vary broadly, in a trait-like manner across individuals (Werth et al., 1997), constant over different nights within a person, influenced by genetic factors, and resistant to experimental perturbations (De Gennaro et al., 2005, 2008; Purcell et al., 2017). Given that spindle frequency is associated with specific anatomical differences in brain structure (Saletin et al., 2013), the measure of spindle frequency may reflect individual differences in functional brain anatomy rather than sleep-specific mechanisms. Thus, participants with faster spindle frequency might generally show larger sleep-dependent consolidation benefits routed in anatomical advantages, rather than one individual’s spindle peak frequency changing with more efficient processing of specific memory content during sleep. Despite the strong effect of spindle peak frequency on memory consolidation found in our meta-analysis, more research is needed to confirm these findings. Further studies are needed to provide a more robust understanding of the relationship between spindle peak frequency and memory consolidation, as has been done for alpha peak frequency regarding its role in cognitive performance (Cecere et al., 2015; Haegens et al., 2014; Mahjoory et al., 2019; Mierau et al., 2017).

The role of spindle power in brain plasticity has been previously shown in research on both animals and humans. In rodents, it has been demonstrated that spindle power is related to the degree of synchrony in neuronal activity, network synchronization, the rate of neural recruitment (Vyazovskiy et al., 2007), and also Ca2+ dendritic activity synchronization (Seibt et al., 2017). After lesions to corticothalamic projections one may expect that the synchrony of spindles is reduced, leading to a decrease in spindle power. Such a power reduction has indeed been observed in patients with unilateral hemispheric stroke (Gottselig et al., 2002) and with damage to the left thalamus (Mensen et al., 2018). Moreover, GABA reuptake inhibitors decrease spindle power and impair sleep-dependent consolidation of sequential motor learning (Feld et al., 2013). Spindle amplitude, which is proportional to spindle power, has also been shown to correlate with the spatial extent of spindle regional recruitment, reflecting a stronger degree of modulation in neuronal firing (Nir et al., 2011). We therefore argue that spindle power, which reflects neural synchrony and the degree of neuronal recruitment, could be directly related to sleep-dependent consolidation.

### 4.4. Exploratory Analyses

The vast majority of sleep and memory research has been conducted in either whole night sleep or nap sleep designs and several studies directly investigated whether the chosen sleep protocol directly affects sleep-memory associations (Cousins et al., 2021; Lo et al., 2014; Mednick et al., 2003; Pöhlchen et al., 2021; Sugawara et al., 2018), especially with regards to the role of sleep spindles in memory consolidation (Cousins et al., 2021; van Schalkwijk et al., 2019). A recent study on procedural memory observed naps and whole nights of sleep may both yield similar memory benefits, suggesting that daytime naps protect memories from deterioration, whereas whole night sleep improves performance, especially in procedural memory (van Schalkwijk et al., 2019). As spindle count and the number of reactivation events may increase with a full night of sleep, and the timing of sleep may have an impact on spindle density (van Schalkwijk et al., 2019), one could expect that nocturnal sleep would result in larger sleep-memory associations as compared to napping. In the current meta-analysis, however, no significant differences between sleep protocols were found, neither across all studies, nor separately in studies reporting declarative or procedural memory. We further confirmed this finding by moderation analyses with sleep duration, showing that sleep duration has no effect on the size of the observed spindle-memory association. Our results align with five previous meta-analyses reporting nearly the same effect size for short and long sleep durations in motor memory (Schmid et al., 2020) and also reporting no moderating effect of sleep protocol in episodic (Hokett et al., 2021) or emotional memory (Schäfer et al., 2020). Similarly, total sleep time was not found to be a significant predictor for an increase in the sleep benefit in episodic memory (Berres and Erdfelder, 2021) and was also not associated with memory and cognition in healthy older adults (Qin et al., 2022). It thus seems that spindles are a general mechanism for memory consolidation, regardless of sleep duration or the timing of sleep.

In the sleep literature, age-related brain alterations have been related to sleep physiology, and sleep spindles, in particular (Fogel et al., 2012, 2014; Martin et al., 2013; Muehlroth et al., 2020, 2019). As older adults also do not show sleep benefits in episodic memory performance (Gui et al., 2017), it is intriguing to ask whether age plays a moderating role in the strength of the association between sleep spindles and memory. In our meta-analyses, however, we did not observe this modulatory effect of age, which aligns with other recent meta-analyses reporting that young and older adults maintain similarly strong sleep-memory relationships (Hokett et al., 2021). This finding is expected, as our sample is relatively young (M = 22.48, age range: 19-37 years), which limits our ability to examine age-related differences for some of the variables of interest. Since sleep studies are predominantly carried out with younger participants (Hokett et al., 2021), future research should more often include older individuals.

### 4.5. Statistical Considerations

In sleep and memory studies, methodological flexibility has often been combined with a shotgun analytical approach, in which a high number of statistical analyses were performed based on several M/EEG scalp locations, spindle types, or spindle measures (see: recent technical review by (Cox and Fell, 2020), with any significant effect interpreted as meaningful. While the studies reviewed here performed on average more than 20 statistical tests, only less than half of the studies used a correction for multiple comparisons. One way to diminish the risk of false positives in high-dimensional brain imaging data, such as multi-channel EEG recordings, is nonparametric permutation testing, which can take into account the spatial or temporal distribution of M/EEG data to reduce the number of comparisons or to adequately control for them (Maris and Oostenveld, 2007). However, only one study reviewed here implemented such cluster-level statistics on the correlation between sleep spindles and behavioral data (Dehnavi et al., 2019). It would be desirable that in the future, more studies looking at spindle-memory associations accounted for multiple comparisons in a similar way.

While any individual researcher has the freedom of defining a statistical threshold and the number of statistical tests they perform, we suggest that the field of sleep and memory research as a whole would benefit if these analytical approaches were chosen prior to statistical testing even before data collection has begun (Cox and Fell, 2020), as implemented in study preregistrations. Preregistrations motivate researchers to formulate hypotheses before seeing the data and can help research transparency and reproducibility by reducing biases (Gorgolewski and Poldrack, 2016; Ioannidis et al., 2014).

Another important statistical consideration is low statistical power because of small sample size (n) that poses a particular threat to the validity of sleep studies. It is known that studies with a small n can lead to an estimation of inflated effect sizes (Schäfer and Schwarz, 2019). In the current meta-analysis, we similarly observed this inflated effect size with smaller sample sizes. Correlating effect size and sample size across studies resulted in a significant association (Supplementary Fig 3), suggesting a clear indication of bias: larger effects result from smaller samples, while smaller effects were observed in the larger samples. We therefore suggest future studies should aim for larger sample sizes and complete an a priori sample size estimation to assure sufficient statistical power (Cordi and Rasch, 2021). This would reduce the risk of obtaining spuriously high effect size measures due to the higher variability in these estimates at a smaller sample size. Since sleep studies have high monetary and time demands, collaborative networks and data-sharing initiatives may also be a promising future avenue for the advancement of sleep science.

Lastly, we observed substantial publication bias in our metaanalysis, similar to other recent meta-analyses of sleep and episodic memory performance (Hokett et al., 2021), complex associative memory processing (Chatburn et al., 2014) or explicit motor sequence learning (Pan and Rickard, 2015; Rickard et al., 2022; but see: Schmid et al., 2020). The results of meta-analyses may overestimate the true effect if publication bias is observed (Ferguson and Brannick, 2012). Several techniques have been developed to correct for publication bias, including trim-and-fill analyses (Duval and Tweedie, 2000a, 2000b) or precision-effect test and precision-effect estimate with standard errors (PET-PEESE; Bartoš et al., 2022). In the current meta-analysis, we employed a moderation analysis testing whether reported effect sizes in published articles were significantly larger than unreported effect sizes. We observed no differences between reported vs. unreported effect sizes across all studies. Unexpectedly, when performing this analysis separately for studies of declarative and procedural memory, we found that reporting status had significant yet opposite effects on the size of the spindle-memory association. While correlation coefficients for declarative memory were higher when reported in the literature, indicating that publication bias led to an overestimation of the spindle-memory association in these studies, for procedural memory, the unreported correlation coefficients showed higher values. This suggests that the spindle-memory association may actually have been underestimated for procedural tasks.

Using our publicly available datasets in OSF (https://osf.io/wu6d7), Ujma (2022) have recently computed the association between sleep spindles and memory by adjusting publication bias using a non-standard multivariate implementation of the above-mentioned PET-PEESE methods. He observed that correcting for publication bias in this way reduces the correlation between sleep spindles and memory function to insignificance. Although this analysis was only performed on all effect size measures we obtained without considering whether general learning ability, memory consolidation, procedural memory consolidation or declarative memory consolidation was investigated, this finding still raises an important question: if spindles were not related to memory consolidation, why do they appear so strongly associated with memory reactivation during sleep (Cairney et al., 2018; Schönauer et al., 2017; Schreiner et al., 2021; Wang et al., 2019)? More importantly, how could they have a causal role in relaying the behavioral effects of targeted memory reactivation (Antony et al., 2018)? We would argue that rather than refraining from studying the role of sleep spindles in human memory because it may not yield significant results, research needs to more directly address the functional relevance of spindles in memory formation. Lastly, given the significant heterogeneity in effect sizes and the low sample sizes reported in the studies included in this meta-analysis, we urge a continued evaluation of true effect sizes and publication bias as the field progresses.

## 5. Limitations

This meta-analysis aimed to assess the strength of the correlation between sleep spindles and memory performance. Given this objective, we have not assessed learning-induced changes in sleep spindles. Therefore, investigating the impact of new learning on subsequent sleep seems to offer a promising direction for future meta-analytic research on sleep spindles and memory consolidation. We further excluded several studies, in particular, studies using manipulations (e.g., targeted memory reactivation, TMR). In TMR, the timing of cue presentation relative to sleep spindles and to the phase of ongoing SOs determines their efficacy in reactivation and memory consolidation (Antony et al., 2019, 2018; Cox et al., 2014). More research is needed to determine how strongly TMR impacts the relationship between sleep oscillatory events and memory performance. A focus on coupling between SO and sleep spindles could provide valuable additional insights into the nature of the relationship between sleep spindles and memory consolidation. However, this question was beyond the scope of the current meta-analysis. While a strength of the current study is a large participant sample and a high number of estimates, most of the included individual experiments had relatively small sample sizes, which might result in an imprecise estimation of the respective effect sizes. Future studies in the field of sleep and memory research should thus aim for larger sample sizes. Lastly, we broadly clustered EEG channels for the moderator analyses into frontal, central, and parietal regions, due to a large variability in recording methods between studies. Using this broad clustering, we found no moderating effect of spindle location on the spindle-memory association. Finer regions of interest, or other topographical approaches would allow investigation of this relationship with better spatial precision. This avenue seems highly promising in light of recent findings that spindle topographies track regional engagement during memory encoding (Petzka et al., 2022). In this regard, it may be fruitful to pursue the question of spindle topography and memory association further depending on different learning tasks.

## 6. Conclusions

In the present study, we examined meta-analytical evidence for an association between sleep spindles and memory performance. We assessed how multiple factors, including memory type, spindle type, different spindle measures, and EEG scalp topography affect this relationship. The results of this metaanalysis have demonstrated a medium but robust association between sleep spindles and memory that was largely independent of spindle type or spindle topography. Collectively, our findings suggest that sleep spindles are involved in learning, and may present a general physiological mechanism for sleep-dependent memory consolidation. Spindle power, in particular, may be a promising predictor of sleep-dependent memory consolidation. We hope future studies can take advantage of our findings to highlight the critical gaps in the understanding of the complex interaction between sleep spindles and memory.

## 7. Data availability statement

All data files, associated codebooks, and analysis scripts are publicly available on the study’s page on the Open Science Framework: (https://osf.io/wu6d7/).

## 8. Acknowledgement

We thank all authors, who shared their data (of the nonsignificant results for the correlation values between spindles and memory) openly with us, which extensively improved the manuscript. We further thank Eduard Stroukov for his help on data assessment and Dorothee Pöhlchen and Nora Roüast for their insightful comments on a previous draft of this manuscript.

## 11. Supplementary Material

**Supplementary Figure 1.**
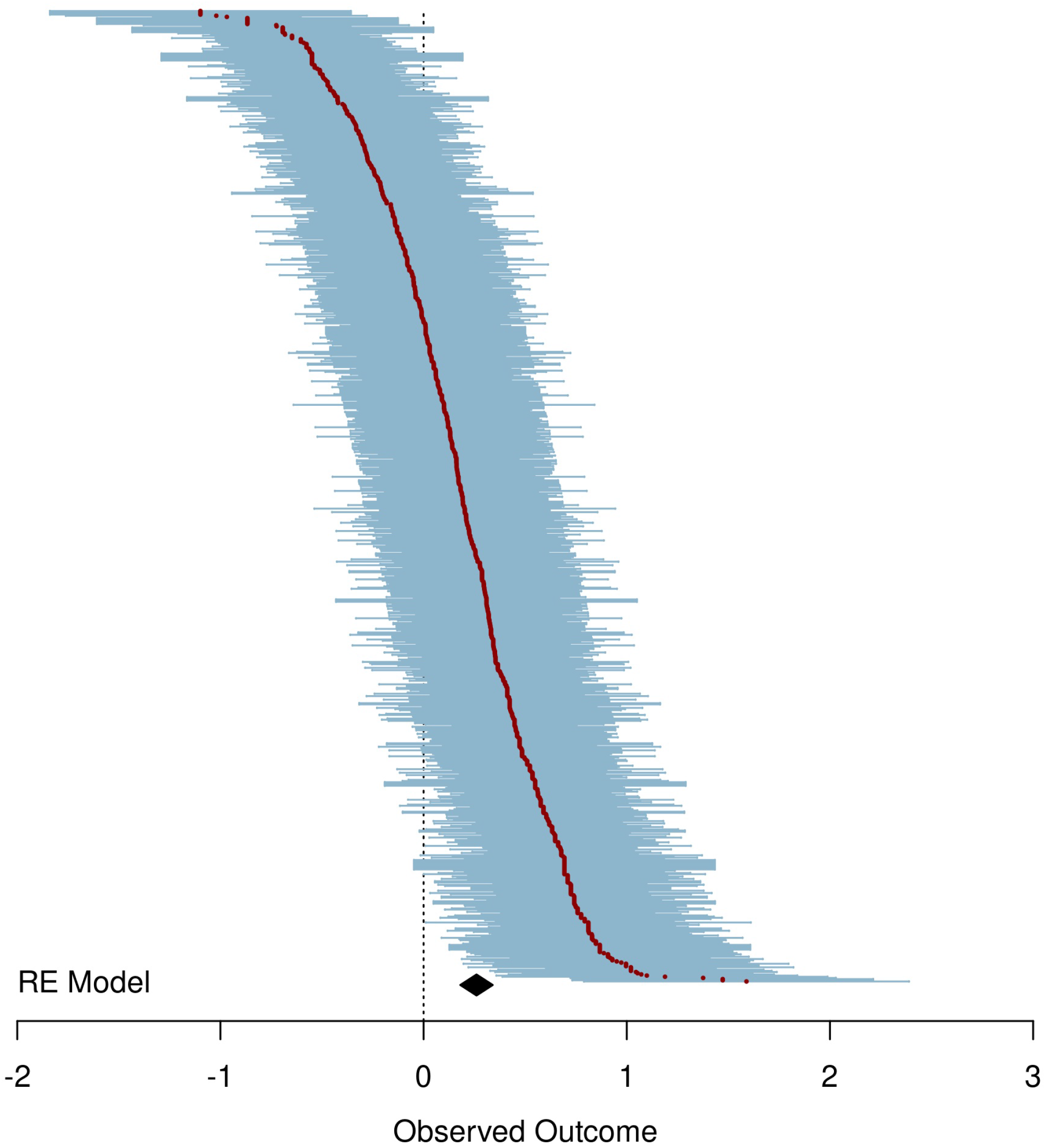
Forest plot for the (r) between levels of sleep spindles and long-term memory, reflecting memory consolidation (k = 1204). Correlations are shown for all effects in the meta-analysis. The size of each dot reflects the weight given to the observed effect during model fitting. The diamond at the bottom shows the meta-analytically weighted mean correlation (with 95% CI, z = 0.240, r = 0.236). The dotted vertical line represents an effect size of 0. Multiple measures were adjusted for dependency using multilevel random effect models (See: **Method section**).

**Supplementary Figure 2.**
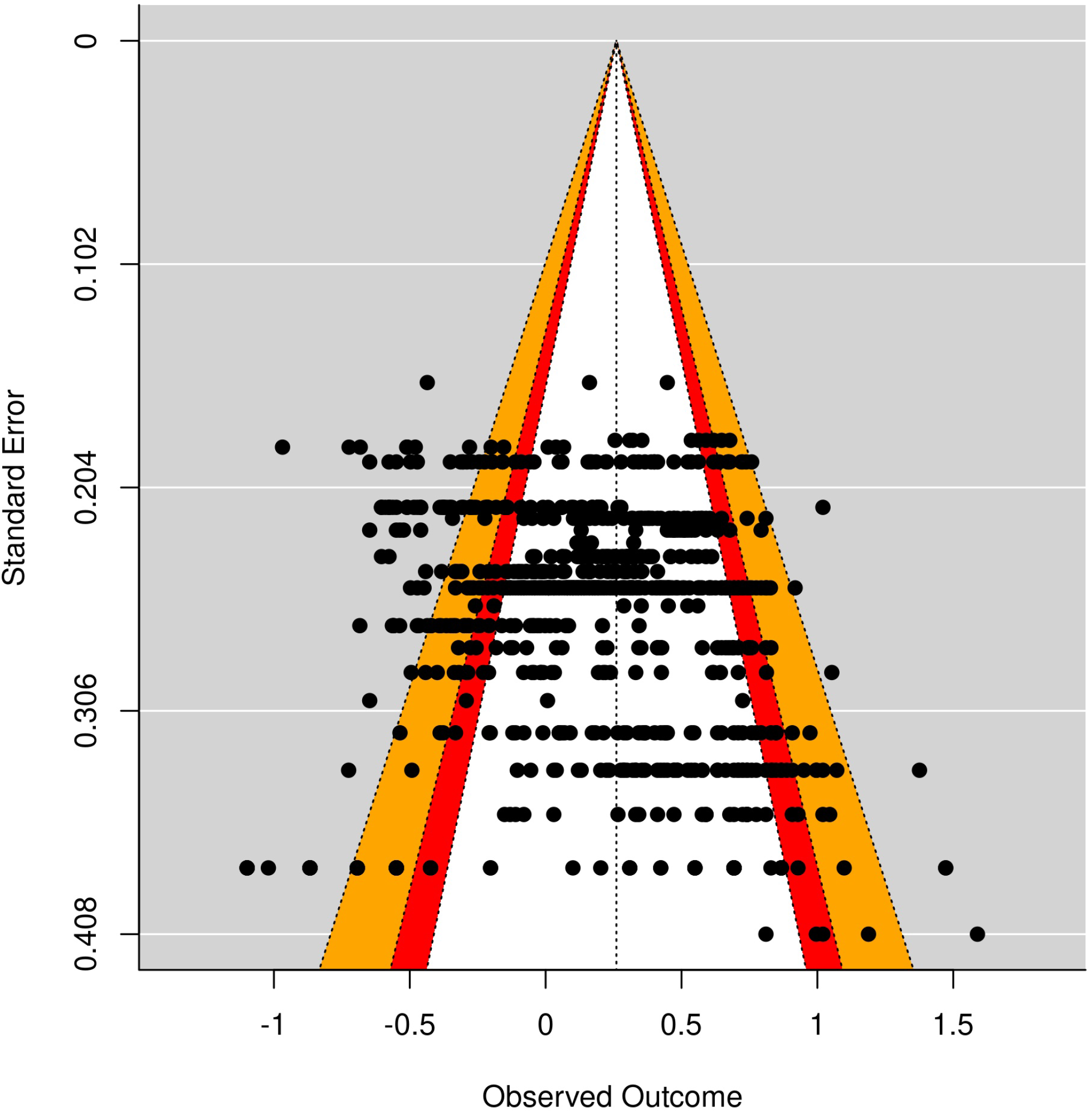
Funnel plot showing the standard errors of effect sizes between sleep spindles and memory consolidation across all studies with 1204 unique associations. The key areas of statistical significance have been superimposed on the funnel plot. While the red zones show effects between *p* = 0.1 and *p* = 0.05, and the orange zones show effects between *p* = 0.05 and *p* = 0.01.

**Supplementary Figure 3.**
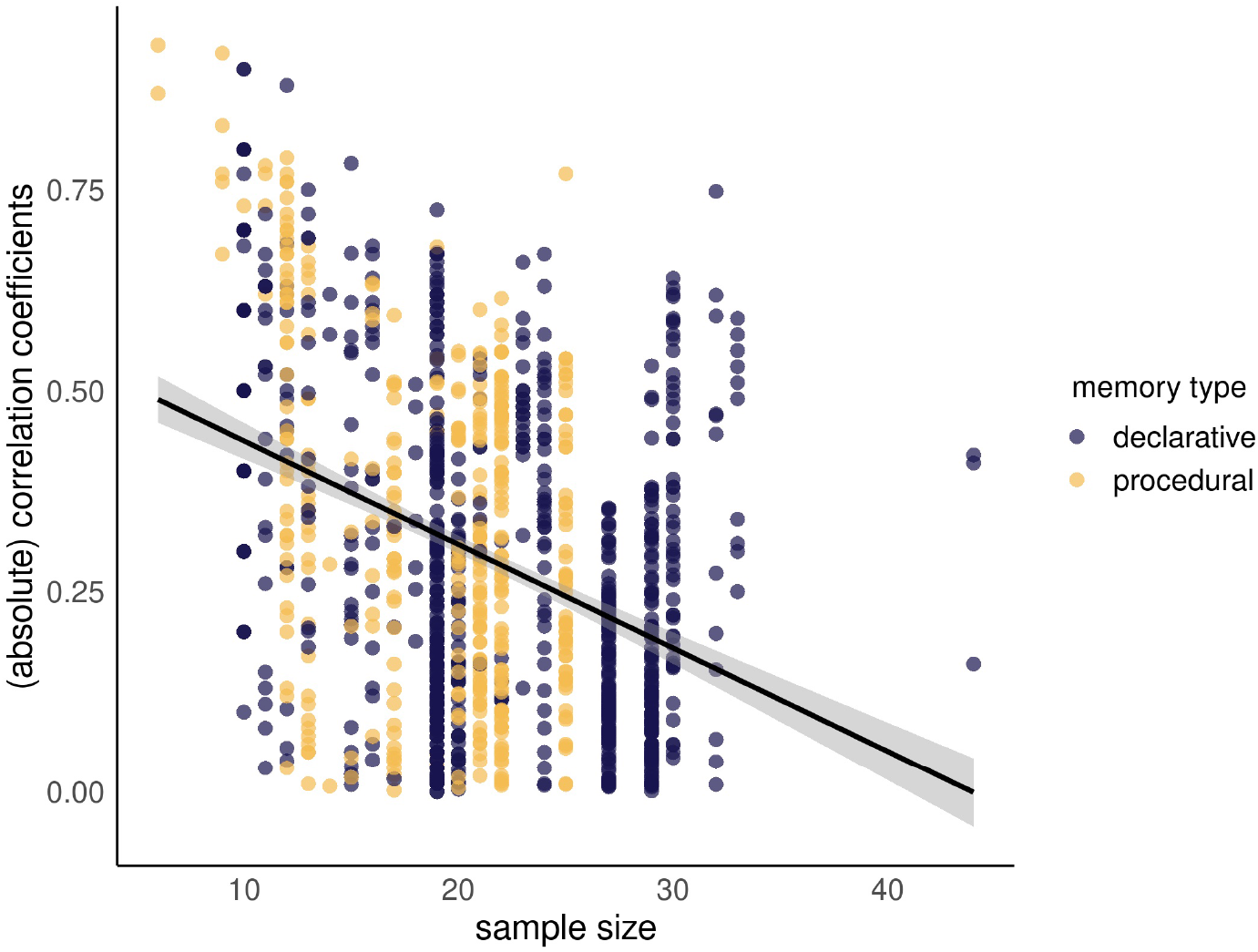
The scatter plot shows the relationship between sample size and effect size. Pearson’s correlation across all studies, reporting memory consolidation resulted in a significant association (r = −0.175, *p* < 0.001, k = 1204) suggesting a clear indication of a publication bias taking the form that smaller effects would result from larger samples and vice versa. Note that to increase the interpretability of the graph, we visualized the correlation plot by excluding one study (Ackermann et al., 2015) with a higher sample size (N = 929 and r = −0.377, *p* < 0.001)).

**Supplementary Table 1.** The number of estimates for each measure of interests across extracted 53 studies with 1424 effect sizes.

| **Variables** | **N** |
| --- | --- |
| total estimates | 1427 |
| learning ability (trait) | 223 |
| memory consolidation (state) | 1204 |
| memory retention | 592 |
| post-sleep | 612 |
| **Memory types** |  |
| declarative memory | 939 |
| procedural memory | 488 |
| hippocampal tasks | 216 |
| non-hippocampal tasks | 272 |
| **Spindle features** |  |
| fast | 610 |
| slow | 348 |
| global | 469 |
| **EEG scalp topography** |  |
| frontal | 428 |
| central | 485 |
| parietal | 275 |
| others (e.g., temporal, occipital) | 239 |
| **Spindle measures** |  |
| amplitude | 164 |
| count | 340 |
| density | 372 |
| duration | 172 |
| frequency | 36 |
| power | 207 |
| spindle activity | 136 |
| **Sleep protocol** |  |
| sleep | 779 |
| nap | 580 |
| others (e.g., half-night, cycle) | 68 |
| **Effect size status** |  |
| Reported | 984 |
| Unreported | 443 |

**Supplementary Table 2.** Post-hoc comparisons for the moderation analysis of spindle measures on spindle-memory associations. *Note that p-values are adjusted according to Bonferroni correction*. ***<0.001,**<0.01,*<0.05, .<0.1

| comparisons | Estimate (difference) | $p_{adj}$ | |
| --- | --- | --- | --- |
| count-amplitude | -0.12 | 0.225 |  |
| density-amplitude | -0.052 | 1 |  |
| duration-amplitude | -0.118 | 0.0145 | * |
| frequency-amplitude | 0.344 | <0.001 | *** |
| power-amplitude | 0.05 | 1 |  |
| SpA-amplitude | -0.133 | 1 |  |
| density-count | 0.068 | 1 |  |
| duration-count | 0.001 | 1 |  |
| frequency-count | 0.464 | <0.001 | *** |
| power-count | 0.169 | 0.961 |  |
| SpA-count | -0.014 | 1 |  |
| duration-density | -0.067 | 0.659 |  |
| frequency-density | 0.396 | <0.001 | *** |
| power-density | 0.102 | 1 |  |
| SpA-density | -0.081 | 1 |  |
| frequency-duration | 0.463 | <0.001 | *** |
| power-duration | 0.168 | 0.827 |  |
| SpA-duration | -0.015 | 1 |  |
| power-frequency | -0.295 | 0.06 | . |
| SpA-frequency | -0.478 | <0.001 | *** |
| SpA-power | -0.183 | 1 |  |

**Supplementary Table 3.** Post-hoc comparisons for the moderation analysis of spindle measures on spindle-memory associations after excluding 2 studies reporting frequency measure (N = 27). *Note that p-values are adjusted according to Bonferroni correction*. *** < 0.001, ** < 0.01, *< 0.05, .< 0.1

| comparisons | Estimate | $p_{adj}$ |
| --- | --- | --- |
| count-amplitude | -0.109 | 0.2964 |
| density-amplitude | -0.046 | 1 |
| duration-amplitude | -0.111 | 0.0225 * |
| power-amplitude | 0.055 | 1 |
| SpA-amplitude | -0.128 | 0.9742 |
| density-count | 0.063 | 1 |
| duration-count | -0.002 | 1 |
| power-count | 0.164 | 0.7887 |
| SpA-count | -0.019 | 1 |
| duration-density | -0.065 | 0.5365 |
| power-density | 0.101 | 1 |
| SpA-density | -0.082 | 1 |
| power-duration | 0.166 | 0.6319 |
| SpA-duration | -0.017 | 1 |
| SpA-power | -0.183 | 0.9867 |

